# CRISPR-Based DNA Imaging in Living Cells Reveals Cell Cycle-Dependent Chromosome Dynamics

**DOI:** 10.1101/195966

**Authors:** Hanhui Ma, Li-Chun Tu, Ardalan Naseri, Yu-Chieh Chung, David Grunwald, Shaojie Zhang, Thoru Pederson

**Author notes:** Co-first authors (Hanhui Ma, Li-Chun Tu).

## Abstract

In contrast to the well-studied condensation and folding of chromosomes during mitosis, their dynamics in interphase are less understood. We developed a sensitive, multicolor system, CRISPR-Sirius, allowing the real-time tracking of the dynamics of chromosomal loci. We tracked loci kilobases to megabases apart and found significant variation in the inter-locus distances of each pair, indicating differing degrees of DNA contortion. We resolved two distinct modes of dynamics of loci: saltatory local movements as well as translational movements of the domain. The magnitude of both of these modes of movements increased from early to late G1, whereas the translational movements were reduced in early S. The local fluctuations decreased slightly in early S and more markedly in mid-late S. These newly observed movements and their cell cycle-dependence are indicative of a hitherto unrecognized compaction-relaxation dynamic of the chromosomal fiber operating concurrently with changes in the extent of observed genomic domain movements.

**IN BRIEF:** Distinct chromosome folding and dynamics during cell cycle progression were dissected by CRISPR-Sirius DNA imaging in living cells.

**HIGHLIGHTS:**

- CRISPR-Sirius allows tracking of pairs of chromosomal loci having kilobase to megabase inter-locus distances
- Pair-wise tracking of loci allows measurement of both local and domain dynamics
- Chromosomal fiber relaxation is positively correlated with local dynamics
- Genomic region size contributes to local and domain movements
- Distinct chromosome dynamics were uncovered during cell cycle progression in interphase

## INTRODUCTION

Proper spatial organization and dynamics of chromosomes at the kilobase to megabase scales are essential for gene regulation and cellular function (Risca et al., 2015). DNA FISH and super-resolution microscopy in fixed cells allow measurement of the distances between loci at tens of nanometers resolution (Boettiger et al., 2016) while chromosome capture identifies DNA loops as well as apparently-autonomous genomic domains and compartments at kilobase resolution by measuring contact probabilities between pairs of loci in a cell population (Lieberman-Aiden et al., 2009; Rao et al., 2014). These methods have revealed a number of chromosome structure features, in a static mode, including folding, compaction and organization in different cellular conditions. Chromatin compaction has been found to be regulated at all scales investigated so far, from the entire chromosome such as the X (Teller et al., 2011), or large intra-chromosomal domains such as the polycomb gene locus (Francis et al., 2004; Boettiger et al., 2016), to the level of single genes (Benabdallah et al., 2017). An inherent limitation of chromosome capture studies, *viz.* that they had been limited to ensemble measurements made on populations of cells, has recently been overcome by refinements that enable singe cell-based analysis (Nagano et al., 2013; Stevens et al., 2017; Flyamer et al., 2017; Nagano et al., 2017).

Because relatively little is known about the inter-locus distances and domain dynamics of interphase chromosomes in living cells, we developed a new system, “CRISPR-Sirius”, which enabled resolution of these genomic features from the kilobase to megabase scale in real time. Positive correlation of maximal spatial distances between loci and their gyration radii of trajectories suggested a connection between local loci pair contortion / chromosomal fiber relaxation and the region’s dynamics. This correlation was further confirmed during cell cycle progression, where contrasting patterns of chromatin local dynamics were observed as cells moved from early G1 to late S.

## RESULTS

### Mining Chromosome-Specific Repeats for CRISPR-based Imaging

CRISPR-Cas9 has been repurposed for tracking chromosomal loci in living cells (Chen et al., 2013) and subsequently orthogonal Cas9s with their cognate guide RNAs were later developed to target and visualize multiple loci in single cells (Ma et al., 2015; Chen et al., 2016). Engineered sgRNA scaffolds have been designed that have increased the spectral range for distinguishing multiple genomic loci (Ma et al., 2016b; Fu et al., 2016; Shao et al., 2016). In our “CRISPRainbow” system (Ma et al., 2016b) six distinct genomic loci were visualized simultaneously in single cells (Ma et al., 2016b). In that study, chromosome-specific repeats containing at least 100 copies of the Cas9 target sites were chosen as loci. To expand this technology, we realized that we would need to know the complete landscape of chromosome-specific tandem repeats, how many families of these chromosome-specific repeats are present on each chromosome, and the copy number of each tandem repeats. We developed a computational search algorithm to mine the human genome repeat landscape specifically for repeats that are restricted to a given chromosome under our defined threshold (see Methods). We found that there are 30 chromosome-specific repeats that have ≥100 copies of Cas9 target sites, 1244 chromosome-specific repeats that have between 20 and 99 copies and 6373 unique repeats that have between 5 and 19 copies. The detailed distribution of chromosome-specific repeats and copy numbers was analyzed and shown in Figure 1. In addition, all candidate target sites with more than 5 copies were deposited in the CRISPRbar web server (http://genome.ucf.edu/CRISPRbar), which can be used to design guide RNA for CRISPR-based imaging. The precise copy number of target sites needed to visualize and track the dynamics of a genomic locus with our technology remains uncertain at present. Nevertheless, the available genomic locations for labeling will expand from those with 30 (≥100 copies) to ones with 1244 (≥20 copies) in the human genome if the sensitivity of labeling can be driven down to 20 copies. The combining of multiple guide RNAs together to reach the total target site threshold of ≥20 copies in defined regions will make more genomic location available, which can be searched on the CRISPRbar which we have developed. Thus it becomes important to generate a more sensitive CRISPR imaging system.

**Figure 1.**
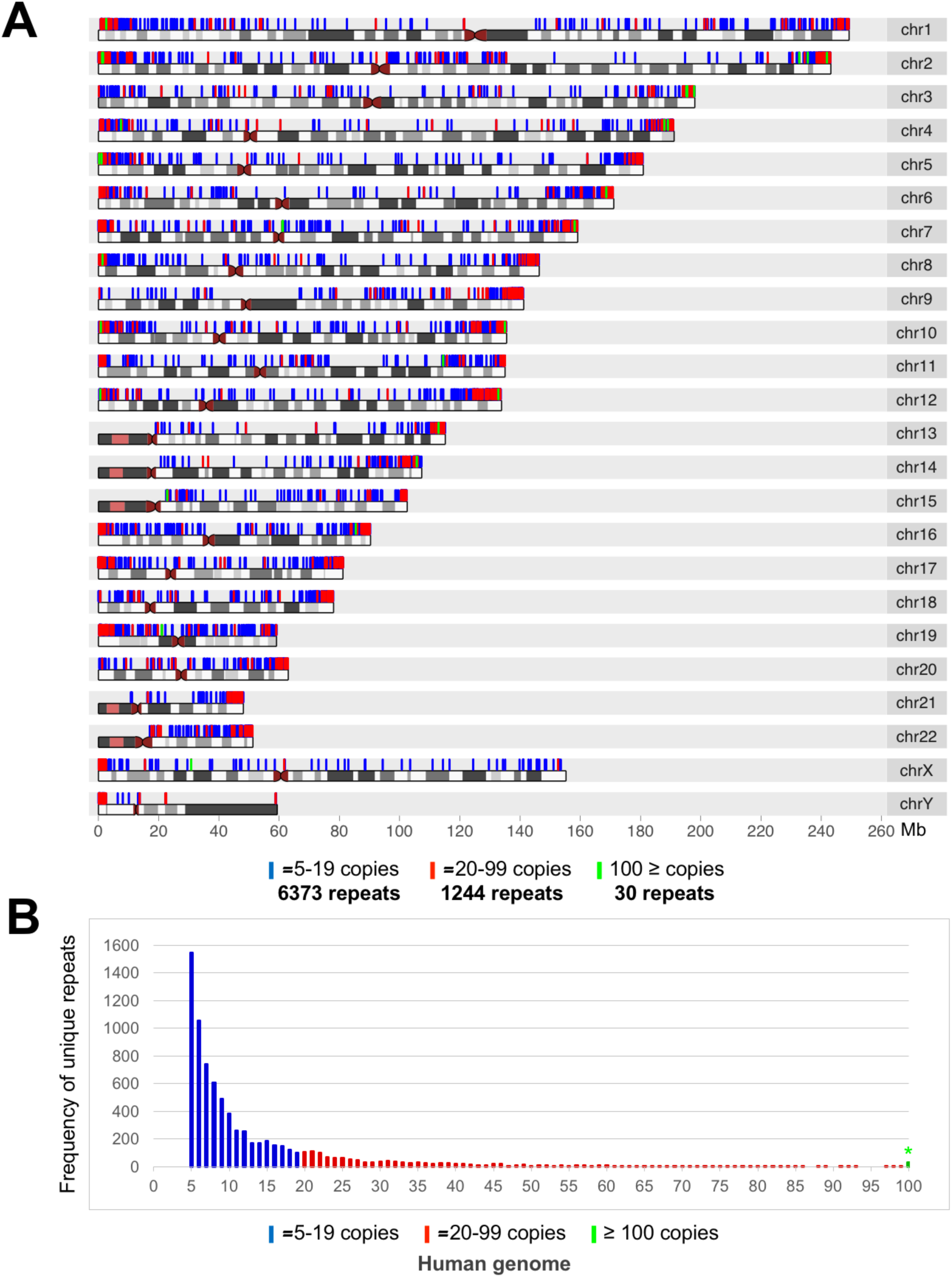
The Distribution of Chromosome-Specific Repeats in the Human Genome. (A) The distribution of chromosome-specific repeats in the human genome is shown. Green represents loci having ≥100 copies of target sites by CRISPR imaging on a repetitive DNA region, red these having 20-99 copies and blue those having 5-19 copies. The copy numbers were defined as the numbers of highest non-overlapping target sites in each repetitive region. (B) Histogram showing the frequency distribution of the chromosome-specific unique repeats as a function of the copy number of the internal core sequence in each: blue: 5-19 copies; red: 20-99 copies; green*: sum of all repeats ≥100 copies.

### CRISPR-Sirius, a Bright, Multicolor DNA Tracking System

An obvious strategy for enhancing CRISPR imaging sensitivity is to amplify the signal strength emanating from each target site. A protein-tagging system termed SunTag was developed to recruit multiple GFPs to a single dCas9 to amplify the fluorescent signals in a single color mode (Tanenbaum et al., 2014). This was a monochromatic advance but, for a polychromatic range, guide RNA-tagging combined with fluorescence enhancement provides a potentially more flexible approach. Placing multiple MS2 RNA hairpins at the 3’-termini of CRISPR sgRNAs has been found to destabilize the RNAs, thus decreasing the efficiency of dCas9:guide RNA complexing and reducing the target labeling (Zalatan et al., 2015; Ma et al., 2016a), although signal amplification was achieved in certain cases (Qin et al., 2017).

Here we developed a method, CRISPR-Sirius, by rational design of sgRNA scaffolds containing octets of either MS2 or PP7 aptamers. Our strategy was to place the aptamer arrays at the end of the repeat:anti-repeat stem of the sgRNA since this region has been shown to accommodate large RNA inserts without perturbing CRISPR targeting (Shechner et al., 2015). In addition, aptamers were linked by three-way junctions to create stable RNA secondary structure (Zuker, 2003). Third, we introduced unique mutations of into individual MS2 and PP7 hairpins to minimize recombination during virus production and limit misfolding of the transcripts (Wu et al., 2015). We named these engineered scaffolds CRISPR Sirius-sgRNA-8XMS2 and Sirius-sgRNA-8XPP7, as diagrammed in Figures 2A and 2B respectively. Figure S1 contains the flowchart for the rational design of the CRISPR Sirius-8XMS2 scaffold.

**Figure 2.**
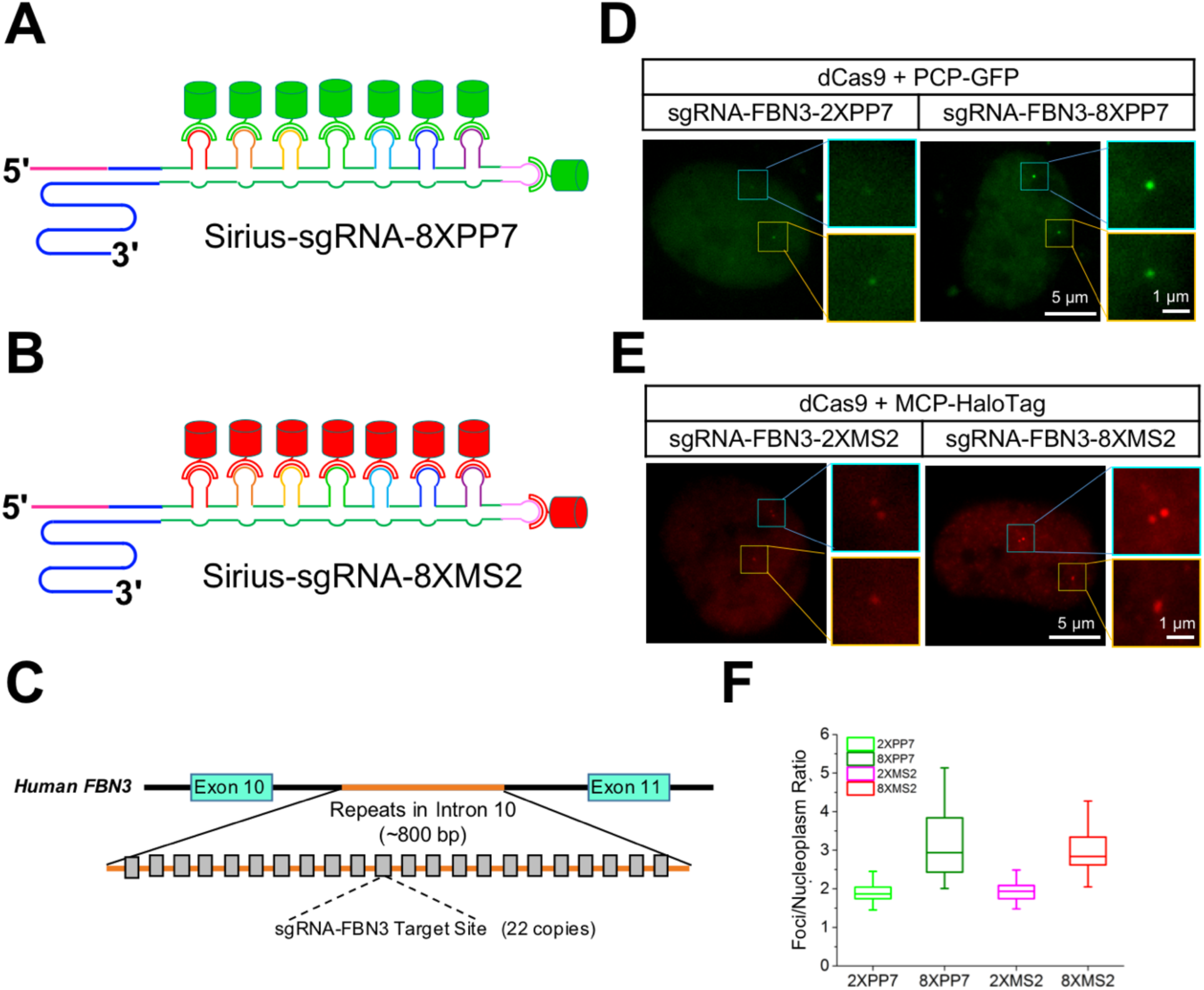
Brightness Increase by Multiplexing RNA Aptamers on the Guide RNA Scaffold. (A-B) Diagrams of CRISPR Sirius-sgRNAs-8XPP7 and Sirius-sgRNA-8XMS2. (C) Schematic of a site in intron 10 of human fibrillin gene *FBN3*, containing 22 copies of target sites. (D-E) Comparison of the foci brightness using CRISPRainbow-sgRNA-FBN3-2XPP7 (left panel in D) or 2XMS2 (left panel in E) and CRISPR-Sirius-sgRNA-FBN3-8XPP7 (right panel in D) or 8XMS2 (right panel in E). The images with the same color were scaled to the same minimal and maximal levels. (F) Quantification of the foci/nucleoplasm intensity ratio with each of the four labeling systems. n=60 loci for each system.

To test the live cell labeling efficiency of low-copy DNA repeats by CRISPR-Sirius, a ∼800 bp region in intron 10 of the human *FBN3* gene was chosen, which consists of 22 tandem copies of the target sites (Figure 2C). The guide RNA complementary this repeat was inserted into either our previously described CRISPRainbow scaffold (sgRNA-FBN3-2XPP7 or 2XMS2) or the new CRISPR-Sirius scaffold (sgRNA-FBN3-8XPP7 or 8XMS2) respectively. The guide RNAs containing PP7 were co-expressed with *S. pyogenes* dCas9 and PCP-GFP, while the those containing MS2 were co-expressed with dCas9 and MCP-HaloTag (Grimm et al., 2015). As seen in Figure 2D and 2E, the target signals from CRISPR-Sirius were considerably brighter than the signals from CRISPRainbow, and the average ratio of foci signal to nuclear background intensity increases as well (Figure 2F). These results showed that CRISPR-Sirius could significantly increase the foci brightness for low-copy repeats, making it possible to visualize many loci with ≥20 copies of target sites. The increase in brightness from CRISPRainbow to CRISPR-Sirius also allows the long-term or successive tracking of multiple loci on a single chromosome.

### CRISPR-Sirius Labeling of Kilobase to Megabase-Spaced Loci on a Single Chromosome

For this study we chose a series of chromosome 19-specific loci for the study of their spatial distance and dynamics. Chromosome 19 spans ∼59 megabases and contains ∼23 genes per megabase, which is the highest gene density of all the human chromosomes (Grimwood J et al., 2004). All the unique repeats containing ≥5 copies of potential Cas9 target sites within 50 kb on the chromosome 19 are shown in Figure S2 or they can be found on web server CRISPRbar. 35 genomic locations each containing ≥20 target site copies were tested for CRISPR-Sirius labeling in U2OS cells (Figure S3). Seven out of these 35 locations-4 intergenic DNA regions (IDRs), 2 intronic regions (TCF3 and FBN3) and 1 pericentromeric region (PR1) on the p-arm of chromosome 19 were chosen to generate six stable cell lines for tracking the dynamics of locus pairs (Figure 3A). We created a dual-guide RNA expression vector for one-step generation of each pair of guide RNAs (Figure S4). IDR3 labeled by CRISPR-Sirius-8XPP7 was used as the common locus in all cell lines while CRISPR-Sirius-8XMS2 was used to label the respective other loci (IDR1, IDR2, TCF3, IDR4, FBN3 and PR1, see Figure 3B). All six pairs of loci were readily visualized in individual cells (Figure 3B). Two foci were detected for IDR1, IDR2, TCF3, IDR3, IDR4 and FBN3 in G1 as expected from the known two copies of chromosome 19 in the U2OS karyotype (Janssen et al., 2013), while only one site was detected for PR1 possibly due to the existence of a deletion mutation in one allele.

**Figure 3.**
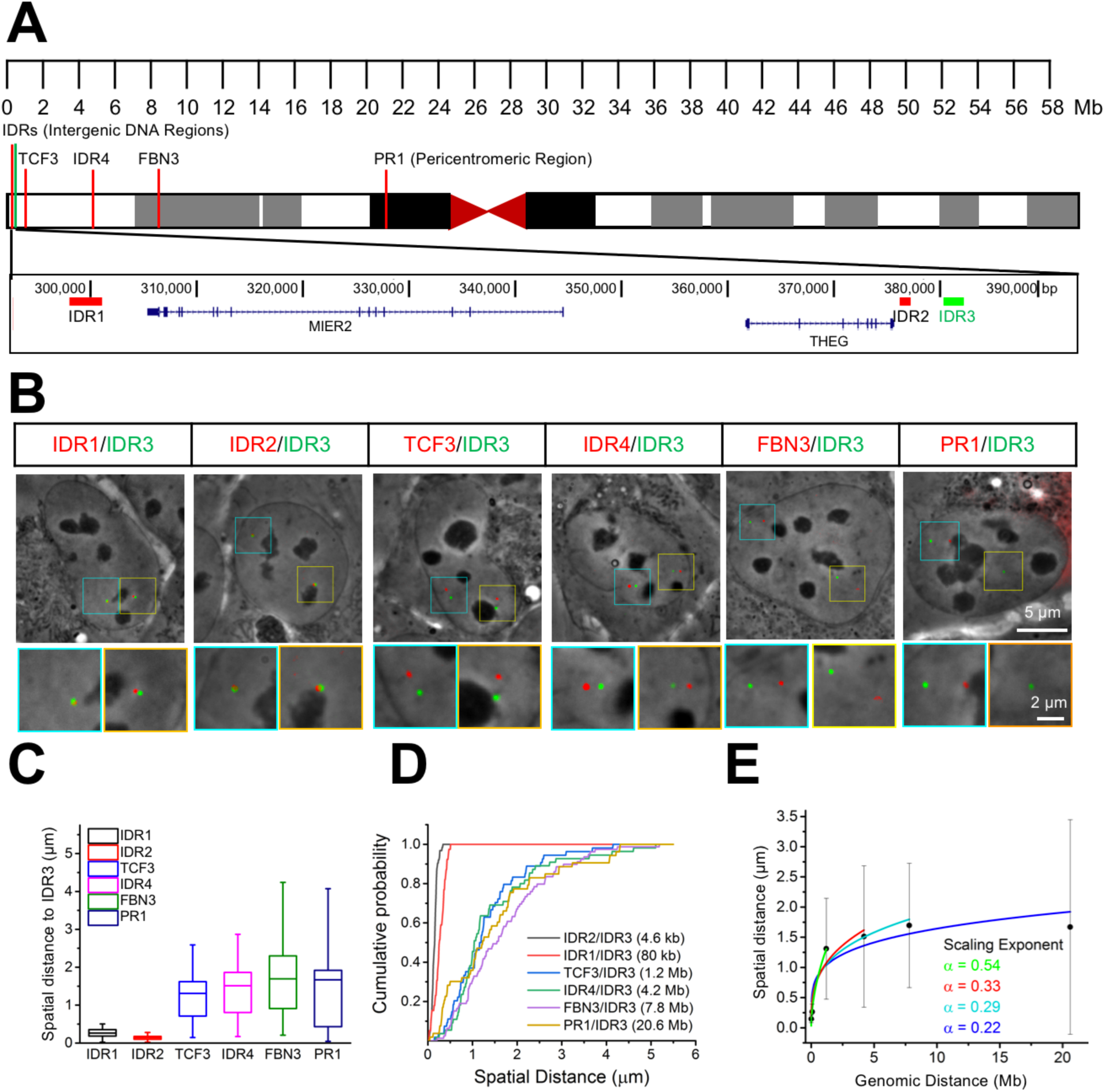
CRISPR-Sirius Labeling of Locus Pairs on the Chromosome 19. (A) Schematic of seven unique loci on human chromosome 19 used in this study consisting of 4 intergenic DNA regions (IDRs), 2 intronic regions (TCF3 and FBN3) and 1 pericentromeric region (PR1). Shown in the lower diagram is an expansion of the region in which IDR1, IDR2 and IDR 3 are so compressed in the upper diagram. (B) Live cell imaging of locus pairs between IDR1/IDR3, IDR2/IDR3, TCF3/IDR3, IDR4/IDR3, FBN3/IDR3 and PR1/IDR3. (C) Plots showing spatial distances between pair-wise combinations of IDR1, IDR2, TCF3, IDR4, FBN3 or PR1 with IDR3. n=40 locus pairs, for IDR1/IDR3, n=62 for IDR2/IDR3, n=54 for TCF3/IDR3, n=56 for IDR4/IDR3, n=78 for FBN3/IDR3 and n=55 for PR1/IDR3. (D) Cumulative probability plot showing the difference of spatial distances distribution of inter-locus distances of each par. (E) Mean spatial distance versus genomic distance for all loci pairs. The scaling exponents α are given by fitting the data to the power law relationship, Mean spatial distance ∞ (genomic distance)^α^.

To quantify the average inter-locus distance of each pair, only those cells in which the pair was on the same focal plane were chosen. As shown in Figure 3C, the observed mean inter-locus distances positively correlated with the DNA physical map, although there was a very high degree of variability among the four most widely spaced pairs of loci (see error bars in Figure 3C), suggesting a diversity of chromosome folding states and inter-locus distances at the kilobase to megabase scales. This is also evident in the cumulative probability plots in Figure 3D. The mean spatial distance between loci when scaled to the DNA physical map has been proposed to plot as a power law with the ∼0.3 exponent expected from an ideal fractal-globule polymer model (Wang et al., 2016). This exponent for the PR1/IDR3 20 Mb domain of chromosome 19 was measured to be 0.22 (Figure 3E), which is nearly identical to the exponent 0.21, estimated in fixed cells by FISH (Wang et al., 2016). The exponents progressively decreased as the inter-locus distance increased while the degree of variation decreased (Figure 3). The degree of variation in the observed exponent for PR1/IDR3 was less for loci spaced at 7.8 kb and 4.2 Mb. These results indicate a positive correlation between the measured inter-locus distances and the known genomic distances within 1 Mb but which deviates when they are further apart.

### Spatial Distance and Dynamics of a Pair of Loci in Single Cells

For the 4.6 kb-spaced IDR2/IDR3 loci (Figure 4A), the measured distance (D_IDR2/IDR3_) ranged from 20 nm to 350 nm in the cell population (Figure 4B). In addition to this extensive variation in the inter-locus distance, we also observed major differences in the dynamic properties of this locus pair. Figures 4C, Movie S1 and Figure 4D, Movie S2 show tracking of IDR2 and IDR3 over time in two different cells. It can be seen that both the range of movements of the site is distinctly greater in cell 2 than cell 1. The trajectory radii for IDR2 (R_IDR2_) and IDR3 (R_IDR3_) were 68 nm and 72 nm respectively in cell 1 (Figure 4C), while they were 153 nm for IDR2 and 180 nm for IDR3 in cell 2 (Figure 4D). The measured distance D_IDR2/IDR3_ ranged from 20-300 nm in cell 2, and 25-110 nm in cell 1 (Figure 4E), which are similar on the minimal distance but remarkably different of maximal distance. To measure the relative local movements of this pair of loci, one (IDR2) was used as a spatial reference and the relative motion of the other (IDR3) (R_IDR2/IDR3_ in Figure 4F) as well as the relative angles were determined as a function of time (θ_R_ in Figure 4J and 4K). The relative trajectory radii of the IDR2 and IDR3 pair was 25 nm in cell 1and 143 nm in cell 2, indicating higher local dynamics of this chromosomal site in the latter (Figure 4G). To measure the movement of the IDR2/IDR3-spanning genomic region itself, the trajectory radius of the centroid (Rc) was calculated (Figure 4H). As shown in Figure 4I, Rc in cell 1 was 69 nm and 151 nm in cell 2, indicating a higher mobility of this domain in the latter.

**Figure 4.**
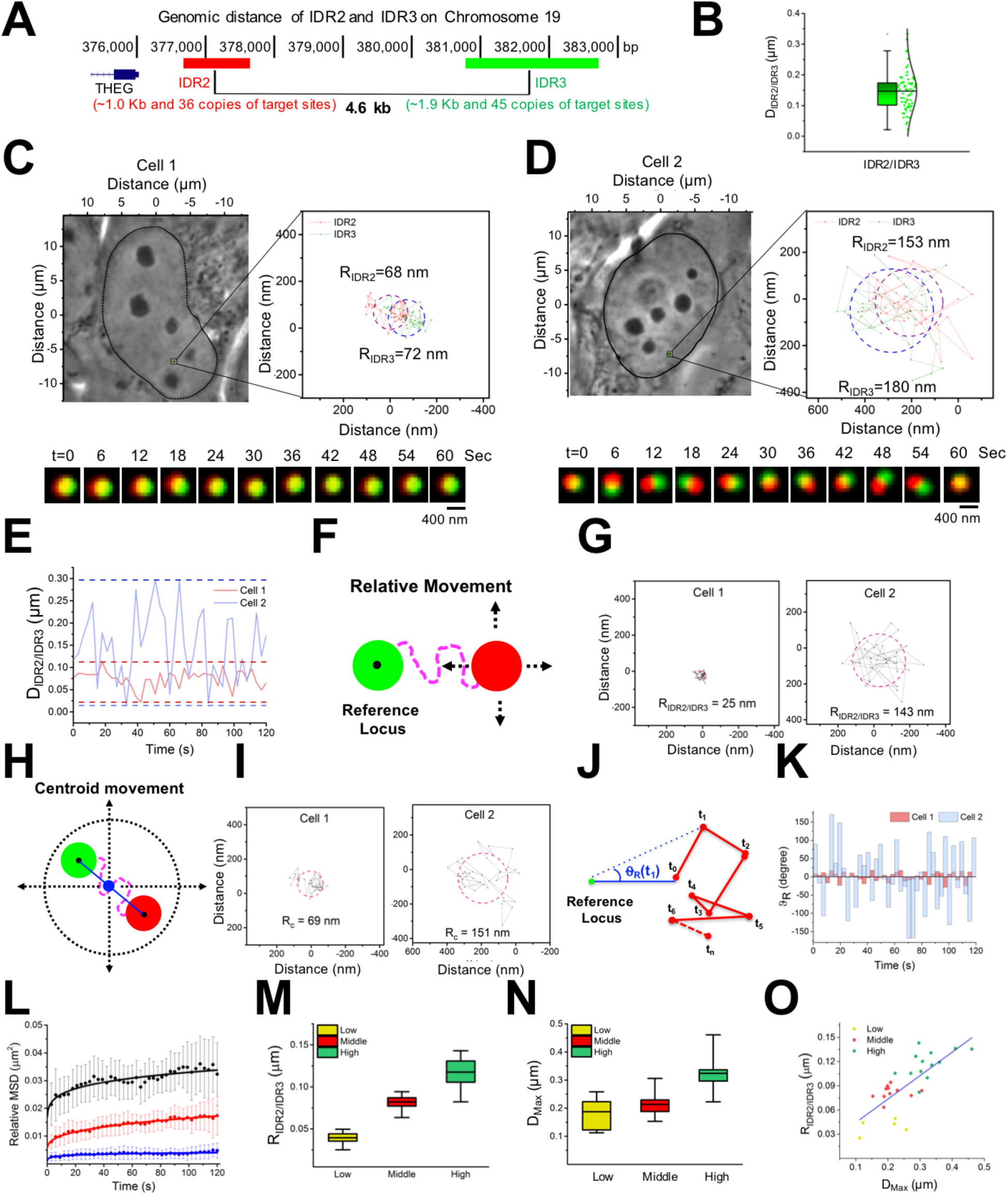
Tracking the Dynamics of a Locus Pair with 4.6 Kilobases Distance. (A) Expanded diagram from Figure 3A showing the IDR2 and IDR3 with their 4.6 kb inter-locus distance. (B) Box and Whisker plot showing the variance of the inter-locus distance between IDR2 and IDR3 from the cell population, n=62 locus pairs. (C-D) Contrasting mobilities of the IDR2/IDR3 locus pair in two cells. Phase contrast images, gyration radius of trajectory (R_IDR2_ and R_IDR3_) and dual-color time lapse images were shown in for each cell. (E) The fluctuating inter-locus distance between IDR2 and IDR3 (D_IDR2/IDR3_) tracked in each cell across time. (F) Schematic of relative movement using the IDR3 locus as a reference point. (G) Relative gyration radii of IDR2 and IDR3 (R_IDR2/IDR3_) in cell 1 and cell 2.(H) Diagram of the centroid movement. (I) Centroid radius of gyration of IDR2 and IDR3 (R_C_) in two cells. (J) Diagram of Relative angles (θ_R_).θ (K) Relative angles of IDR2 and IDR3 (θ_R_) in cell 1 and cell 2 over time.(L) Relative MSD of IDR2 and IDR3 pair was classified into low, middle and high MSD groups. n=13 trajectories, for high, n=9 for middle and n=5 for low MSD. (M-N) Box-and-Whisker plots of relative radius of gyration (R_IDR2/IDR3_) and maximal inter-locus distance (D_Max_) of IDR2 and IDR3 in the low, middle and high MSD groups. (O) Maximal spatial distance (D_Max_) scaled to relative radius of gyration (R_IDR2/IDR3_) of IDR2 and IDR3.

To further understand the relation between the inter-locus distance and the local dynamics, we calculated the mean square displacement (MSD), maximal spatial distance (D_Max_), and relative trajectory radii (R_IDR2/IDR3_). The MSD measures the amount of space the particle explores and when plotted vs. time can reveal whether a particle is in random Brownian motion or subdiffusion in a space (Dion et al, 2013). As can be seen in Figure 4L, the relative MSDs of IDR2/IDR3 fall into three populations. The low population plateaued at ∼0.005 µm^2^, the middle one at ∼0.015 µm^2^ and the high MSD one at ∼0.030 µm^2^. As shown in Figure 4M, the trajectory radii increased from the low to high MSD populations, and the maximal spatial distance (D_Max_), represents the contortion of the locus pair or compaction/relaxation of the chromosomal fiber in this kilobases’ domain also increased from the low to high MSD populations (Figure 4N). The positive correlation of maximal spatial distance and trajectory radii is shown in Figure 4O. These results indicate that the compactness of the chromosomal fiber is closely related to its local mobility.

### Distinct Local and Domain Dynamics during Cell Cycle Progression

A likely basis for the cell-to-cell variation in the dynamic parameters we have observed in these randomly growing cell populations could be the interphase cell cycle stage. In order to address this, we constructed U2OS-derived stable cell lines expressing both dCas9 and the suitable colored guide RNAs to label IDR2 and IDR3 (see Methods), synchronized the cells, and tracked inter-locus and domain movements at different cell cycle stages (Figure 5A). As shown in Figure 5B, the MSD curve of the pair of loci was low in early G1, maximum in late G1, decreased in early S and then declined to a low level again in mid-late S. The average radii of trajectories between the two loci were calculated (Table S1) and are plotted in Figure 5C: 4.04 ± 1.25 × 10^-2^ µm in early G1, 1.09 ± 0.26 × 10^-1^ µm in late G1, 9.35 ± 2.57 × 10^-2^ µm in early S and 5.57 ± 1.51 × 10^-2^ µm in mid-late S. As shown in Figure 5D, the centroid MSD was low in early G1, maximum in late G1, then declining in early S and reaching a minimum in mid-late S. The average centroid radii of trajectory were 8.41 ± 2.17 × 10^-2^ µm in early G1, 1.29± 0.47 × 10^-1^ µm in late G1, 8.79 ± 2.64 × 10^-2^ µm in early S and 5.73 ± 1.13 × 10^-2^ µm in mid-late S (Table S1 and Figure 5E). As shown in Figure 5F, the maximal inter-locus distances underwent similar changes indicating the decompaction of this chromosome region from early G1 to late G1 and then condensation during S phase, which is consistent with results from single cell Hi-C analysis (Nagano et al., 2017) and live cell chromosome accessibility measurements (Pederson and Robbins, 1972). The distribution of relative angles is narrower in early G1 and mid-late S than late G1 and early S (Figure 5G), further supporting the changes in chromosome mobility. These results indicate intrinsic constraints such as folding (Nagano et al., 2017) could be the determinant for local dynamics and external constraints such as the local environment (Ou et al., 2017) might be essential for chromosomal domain dynamics.

**Figure 5.**
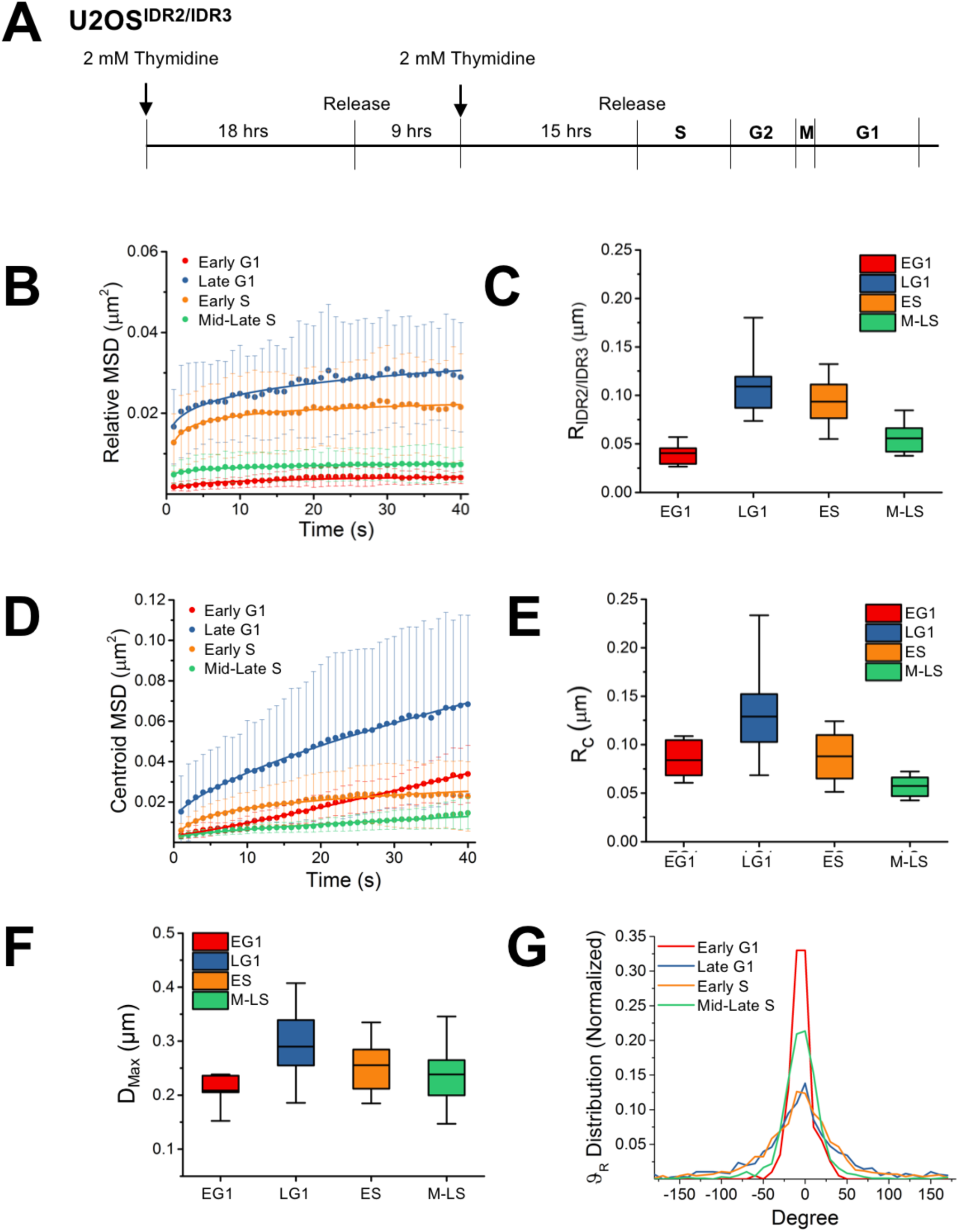
Chromosome Dynamics during Cell Cycle Progression in Interphase. (A) Diagram of cell cycle synchronization for the U2OS^IDR2/IDR3^ cell lines. (B-C) Relative and centroid MSD plot of IDR2/IDR3 in early G1, late G1, early S and mid-late S respectively. n=5 trajectories for early G1, n=19 for late G1, n=21 for early S and n=24 for mid-late S. (D-E) Relative radius (R_IDR2/IDR3_) and centroid radius (R_C_) of IDR2/IDR3 over time in early G1, late G1, early S and mid-late S respectively. (F) Maximal inter-locus distance (D_Max_) of IDR2/IDR3 in early G1, late G1, early S and mid-late S respectively. (G) Relative angle (θ_R_) distribution of IDR2/IDR3 in early G1, late G1, early S and mid-late S respectively.

### The Effect of Genomic Domain Size on the Local and Domain Dynamics

The interphase chromosomes are folded in 3D with a hierarchy of architectures at different length scales from sub-kilobase to megabases (Risca VI et al., 2015). Less is known of the higher-order structure and dynamics beyond nucleosomes at scales such as few to tens of kilobases (*e.g.* regulatory elements), hundreds of kilobases (*e.g.* topologically associating domain (TAD)) from chromosome capture methods (Dixon et al., 2012; Nora et al., 2012) and ∼one megabase chromosomal domains (CDs) visualized by microscopy (Cremer T et al., 2001). We choose four pairs of loci that are spaced difference distances from IDR3 on the DNA physical map by 4.6 kb (IDR2), 80 kb (IDR1), 1.2 Mb (TCF3) and 4.2 Mb (IDR4) (Figure 6A). We then tracked their movements relative to each other and thereby explore how inter-locus distances contributes to the local and domain dynamics of these chromosomal sites. Late G1 phase was chosen for this comparison as substantially local (relative) movement and domain (centroid) movement of the 4.6 kb IDR2/IDR3 pair was observed in this cell cycle stage (Figure 5B-5E).As shown in Figure 6B, the relative MSD for the 4.6 kb pair (IDR2/IDR3) plateaued at ∼0.03µm^2^, the 80 kb pair (IDR1/IDR3) at ∼0.06 µm^2^, while the 1.2 Mb pair (TCF3/IDR3) and 4.2 Mb pair (IDR4/IDR3) MSD curves reached ∼0.14 µm^2^ without plateauing. The average radii of trajectories increased from 1.12± 0. 22 × 10^-1^ µm for the 4.6 kb pair (IDR2/IDR3), or 1.45 ± 0.18 × 10^-1^ µm for the 80 kb pair (IDR1/IDR3) to 2.09 ± 0.51 × 10^-1^ µm for the 1.2 Mb pair (TCF3/IDR3), and no further increase for the 4.2 Mb pair (IDR4/IDR3) (2.09 ± 0.46 × 10^-1^ µm) (Figure 6C and Table S1). These results show that the trajectory radii of relative dynamics expand along with the increase of genomic distance within one megabase but deviate when further apart. When the centroid MSDs were measured however, an opposite pattern emerged for their relationship with genomic distance (Figure 6D and 6E). This contrasting relationship of the centroid MSD vs. the trajectory radius to genomic distance can be clearly seen in the plots in Figure 6F. The distribution of relative angles (Figure 6G) further supports the differences in the mobility of pairs of loci separated by different scales of genomic distances.

**Figure 6.**
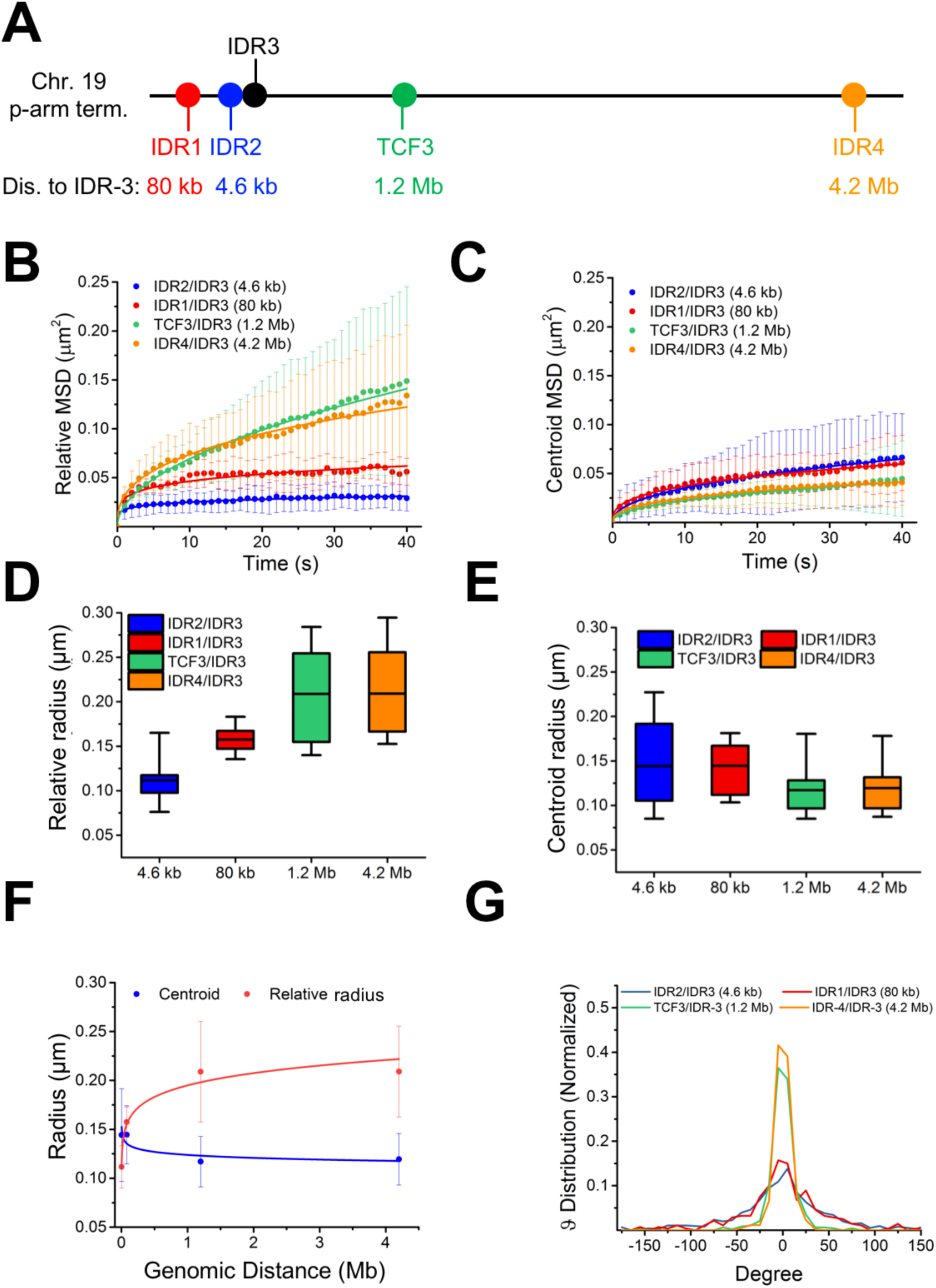
Dynamics of Locus Pairs with Kilobase to Megabase Distances. (A) Diagram of loci on the p-arm of chromosome 19 with distances from IDR3 of 4.6 kb (IDR2), 80 kb (IDR1), 1.2 Mb (TCF3), 4.2 Mb (IDR4). (B-C) Relative and centroid MSD plot of loci pairs from IDR2/IDR3 (4.6 kb), IDR1/IDR3 (80 kb), TCF3/IDR3 (1.2 Mb) to IDR4/IDR3 (4.2 Mb). n=19 trajectories for IDR2/IDR3, n=7 for IDR1/IDR3 pair, n=12 for TCF3/IDR3 and n=11 for IDR4/IDR3. (D-E) Relative radii and centroid radii of the indicated locus pairs. (F) Genomic distance scaled to gyration radius of trajectory. (G) Relative angle (θ_R_) distribution of the indicated locus pairs.

## DISCUSSION

Our perspectives of genomes have transitioned from linear sequences to a hierarchical organization of the chromosomes and their domains in the nucleus (Gibcus et al., 2013; Rao et al., 2014). However, little is understood about interphase chromosome dynamics at the kilobase to megabase scales. Here we applied CRISPR-based multi-color imaging technology to uncover the spatial and temporal features of chromosome dynamics by measuring the spatial distance and mobility of pairs of intra-chromosomal loci in single living cells. We chose pairs of loci ranging from kilobases to megabases apart on chromosome 19. The inter-locus distances of each given pair showed high cell-to-cell variability, in agreement with previous FISH results (Finn et al., 2017a; Finn et al., 2017b; Giorgetti et al., 2016; Fudenberg et al., 2017) but in our case in the live cell mode. To assess local chromosome dynamics during cell cycle progression, we analyzed a pair of loci spaced by 4.6 kb from the center-to-center between the regions labeled by the two CRISPR probes. This region was estimated to include ∼20 nucleosomes and span ∼600 nm at maximum elongation, based on the beads-on-a-string model (Szerlong et al., 2011). There are remarkable differences of spatial distances (ranging from 20-350 nm) and the gyration radii of trajectory (from 20-150 nm) among different cells, indicating different degrees of folding, compaction and mobility of this genomic region. Our results also showed that a high degree of chromosomal fiber relaxation (D_Max_ in Figure O) is accompanied by high local dynamics. The maximal spatial distance of the 4.6 kb loci pair was observed to be up to 450 nm (Figure 4N), which is two thirds of the expected stretched length of ∼600 nm. This suggests that this region is likely composed of irregularly folded 10-nm fibers, rather than a higher order structure such as 30-nm chromatin fibers. State-of-the-art electron microscopy (Ou et al., 2017) of this region in higher resolution will be needed to confirm whether there is a higher order structure. We further demonstrated that the chromosome’s local relaxation and dynamics are cell cycle-dependent with low local mobility at early G1 and mid-late S, and high at late G1 and early S (Figure 7). These observations deviate from 5C and Hi-C data of genome organization during the cell cycle, which showed that chromosome domains such as TADs quickly reform during early G1 and remain during the rest of the cell cycle before the mitosis (Naumova et al., 2013). However recent single cell Hi-C data across the cell cycle showed that condensation of intra-TADs remarkably increased at mid-late S phase (Nagano et al., 2017), which is consistent with our findings of low local chromosomal fiber relaxation and dynamics in mid-late S phase, although direct comparisons of Hi-C data and live cell imaging-tracking studies are presently somewhat challenging.

**Figure 7.**
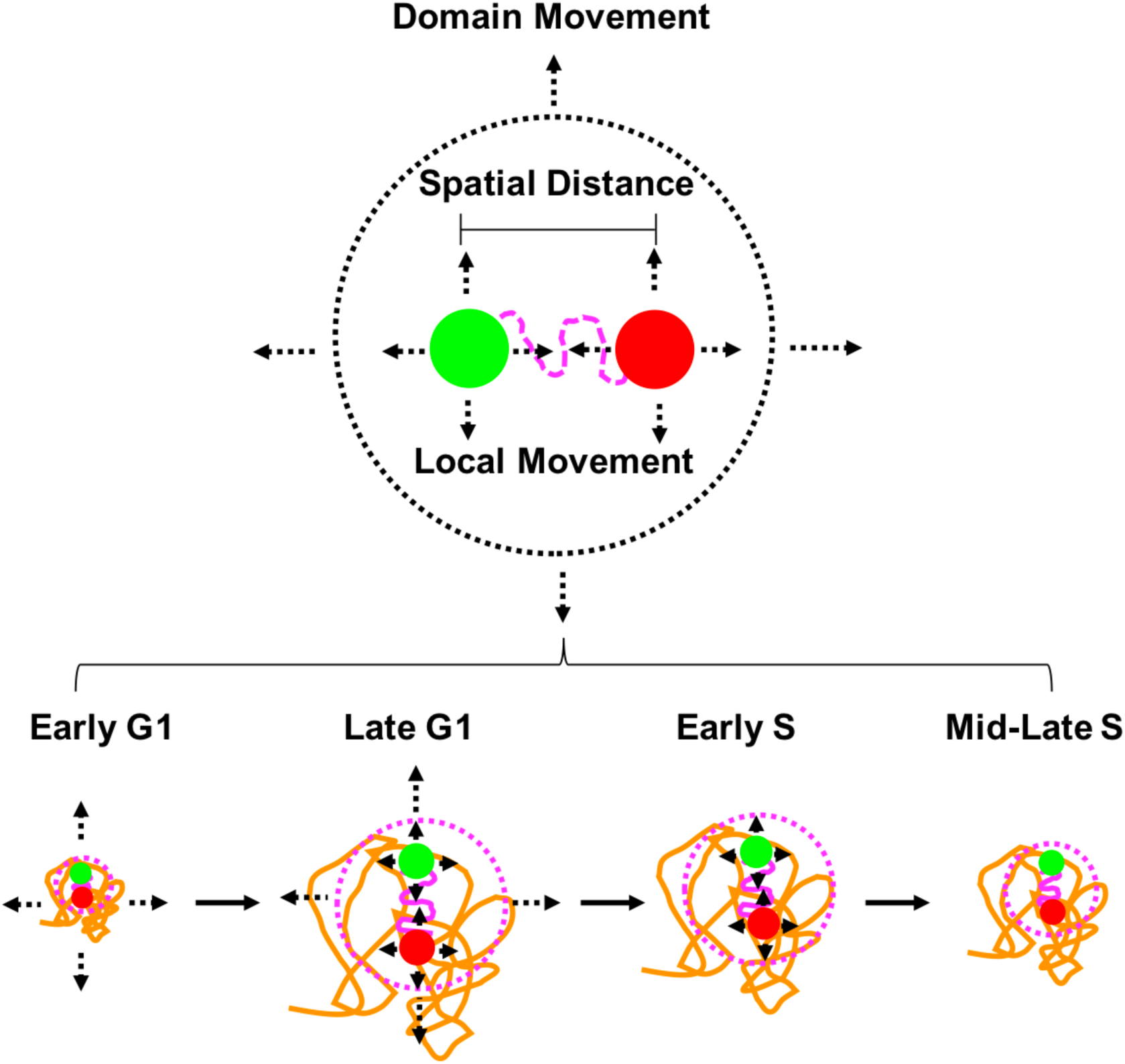
Cell Cycle-Dependent Chromosome Dynamics. Upper: spatial distances and two distinct dynamic modes (local movement and domain movement) can be measured. Lower: Distinct chromosomal fiber relaxation and dynamics were shown during cell cycle progression in interphase.

It is still unclear what determines chromosomal fiber relaxation and dynamics during cell cycle progression or other biological processes. One reason could be the combination of chromatin intrinsic properties and external constraints from the local environment. Here we found that chromosomal fiber relaxation is positively correlated to local dynamics during cell cycle progression. Such compaction changes have also been seen during the epigenetic regulation such as X-chromosome inactivation (Teller et al., 2011), formation of polycomb domains (Francis et al., 2004; Boettiger et al., 2016) and modification of single genes (Benabdallah et al., 2017). Recent electron microscopy has revealed that both euchromatin and heterochromatin contain 5-24 nm fibers, but that their compactness is considerably different (Ou et al., 2017) suggesting that the cooperation between chromosomal fiber relaxation and dynamics could possibly be involved in the transcriptional regulation. In combination with existing protein or RNA labeling techniques (Stasevich et al., 2014; Nelles et al., 2016), our CRISPR-based live cell DNA tracking of genomic loci pairs could be applied to studies of the transcription kinetics in different epigenetic states as well as DNA repair, translocation or recombination (Hauer et al., 2017; Amitai et al., 2017).

## STAR * METHODS

### METHODS DETAILS

- Design of RNA Scaffolds for CRISPR-Sirius
- Plasmid Construction
- Cell Culture, Transfection and Cell Cycle Synchronization
- Lentivirus Production and Transduction
- Flow Cytometry and Stable Cell Line Selection
- Fluorescence Microscopy

### QUANTIFICATION AND STATISTICAL ANALYSIS

- Chromosome-Specific Repeats for Chromosome 19
- Imaging Processing from Pairs of loci Tracking
- Quantification of Distance and Movement from Pairs of loci

## SUPPLEMENTAL INFORMATION

Supplemental Information includes four figures, one table and two movies.

## AUTHOR CONTRIBUTIONS

H.M. L-C.T. and T.P. conceived the project and designed the CRISPR-Sirius system; A.N. and S.Z. designed the Sirius sgRNA scaffold. A.N. and S.Z. performed data mining of chromosome-specific repeats in the human genome; H.M. and L-C.T. designed and performed experiments; Y-C.C. and L-C.T. performed imaging processing and quantification analysis; H.M., L-C.T., Y-C.C., D.G. S.Z. and T.P. interpreted data; H.M. and T.P. wrote the paper with input from all the authors.

## ACKNOWLEDGEMENTS

HaloTag JF-549 was a gift from the laboratory of Luke Lavis lab at Janelia Research Campus, Howard Hughes Medical Institute. We are grateful to Jon Goguen of UMass Medical School for microscopy support, and thank Magda Kordon, Ying Feng and Yang Zhao in the Pederson lab for the help in some of the imaging and data processing. We also thank Aviva Joseph for providing the Heat Stable Antigen (HSA) labeling assay. This work was supported in part by the Vitold Arnett Professorship Fund to T.P., NIH grant U01 DA-040588 to P. Kaufman, J. Dekker and T.P., NIH grant R01 GM102515 to S.Z., NIH grant U01 EB021238 to D.G. and NIH grant (K99 GM126810) to L-C.T. We are also very grateful to Job Dekker, Erik Sontheimer and Scot Wolfe at UMass Medical School for critical comments on the manuscript and strong support during the study.

---

**Table S1.** Biophysical Parameters Extracted from Loci Pair Trajectories, Related to Figure 4, Figure 5 and Figure 6.

| MSD | Curve | Diffusion Constant ($\mu\text{m}^2/\text{sec}^\beta$) | Exponent ($\beta$) | Trajectory Radius ($\mu\text{m}$) | n* |
| --- | --- | --- | --- | --- | --- |
| Figure 4L | High Level | $(4.76 \pm 0.21) \times 10^{-3}$ | $(1.19 \pm 0.11) \times 10^{-1}$ | $(1.18 \pm 0.17) \times 10^{-1}$ | 13 |
| Figure 4L | Middle Level | $(1.55 \pm 0.04) \times 10^{-3}$ | $(2.09 \pm 0.07) \times 10^{-1}$ | $(8.19 \pm 0.82) \times 10^{-2}$ | 11 |
| Figure 4L | Low Level | $(5.28 \pm 0.32) \times 10^{-4}$ | $(1.37 \pm 0.15) \times 10^{-1}$ | $(3.94 \pm 0.94) \times 10^{-2}$ | 5 |
| Figure 5D | EG1_Centroid | $(4.35 \pm 0.33) \times 10^{-4}$ | $(7.82 \pm 0.26) \times 10^{-1}$ | $(8.41 \pm 2.17) \times 10^{-2}$ | 5 |
| Figure 5B | EG1_Relative | $(4.10 \pm 0.19) \times 10^{-4}$ | $(2.68 \pm 0.15) \times 10^{-1}$ | $(4.04 \pm 1.25) \times 10^{-2}$ | 5 |
| Figure 5D | LG1_Centroid | $(3.17 \pm 0.08) \times 10^{-3}$ | $(4.51 \pm 0.09) \times 10^{-1}$ | $(1.29 \pm 0.47) \times 10^{-1}$ | 19 |
| Figure 5B | LG1_Relative | $(4.31 \pm 0.08) \times 10^{-3}$ | $(1.55 \pm 0.06) \times 10^{-1}$ | $(1.09 \pm 0.26) \times 10^{-1}$ | 19 |
| Figure 5D | ES_Centroid | $(1.77 \pm 0.05) \times 10^{-3}$ | $(3.59 \pm 0.10) \times 10^{-1}$ | $(8.79 \pm 2.64) \times 10^{-2}$ | 21 |
| Figure 5B | ES_Relative | $(3.53 \pm 0.07) \times 10^{-3}$ | $(1.30 \pm 0.06) \times 10^{-1}$ | $(9.35 \pm 2.57) \times 10^{-2}$ | 21 |
| Figure 5D | M-LS_Centroid | $(6.60 \pm 0.25) \times 10^{-4}$ | $(4.09 \pm 0.14) \times 10^{-1}$ | $(5.73 \pm 1.13) \times 10^{-2}$ | 24 |
| Figure 5B | M-LS_Relative | $(1.28 \pm 0.02) \times 10^{-3}$ | $(1.07 \pm 0.04) \times 10^{-1}$ | $(5.57 \pm 1.51) \times 10^{-2}$ | 24 |
| Figure 6B | 4.6 kb_Relative | $(4.36 \pm 0.08) \times 10^{-3}$ | $(1.57 \pm 0.06) \times 10^{-1}$ | $(1.12 \pm 0.22) \times 10^{-1}$ | 19 |
| Figure 6B | 80 kb_Relative | $(7.19 \pm 0.17) \times 10^{-3}$ | $(2.08 \pm 0.09) \times 10^{-1}$ | $(1.57 \pm 0.15) \times 10^{-1}$ | 7 |
| Figure 6B | 1.2 Mb_Relative | $(5.33 \pm 0.07) \times 10^{-3}$ | $(5.12 \pm 0.05) \times 10^{-1}$ | $(2.09 \pm 0.51) \times 10^{-1}$ | 12 |
| Figure 6B | 4.2 Mb_Relative | $(7.95 \pm 0.13) \times 10^{-3}$ | $(3.65 \pm 0.06) \times 10^{-1}$ | $(2.09 \pm 0.46) \times 10^{-1}$ | 11 |
| Figure 6C | 4.6 kb_Centroid | $(3.17 \pm 0.08) \times 10^{-3}$ | $(4.43 \pm 0.08) \times 10^{-1}$ | $(1.44 \pm 0.47) \times 10^{-1}$ | 19 |
| Figure 6C | 80 kb_Centroid | $(4.25 \pm 0.11) \times 10^{-3}$ | $(3.41 \pm 0.08) \times 10^{-1}$ | $(1.45 \pm 0.30) \times 10^{-1}$ | 7 |
| Figure 6C | 1.2Mb_Centroid | $(1.98 \pm 0.02) \times 10^{-3}$ | $(4.52 \pm 0.05) \times 10^{-1}$ | $(1.17 \pm 0.26) \times 10^{-1}$ | 12 |
| Figure 6C | 4.2Mb_Centroid | $(2.64 \pm 0.03) \times 10^{-3}$ | $(3.71 \pm 0.05) \times 10^{-1}$ | $(1.19 \pm 0.26) \times 10^{-1}$ | 11 |

\*Number of trajectory

**Movie S1. Tracking of IDR2/IDR3 Loci Pair in Cell 1, Related to Figure 4**

Images were cropped to 150x150 pixels and each video includes 41 frames (a total time of 2 minutes). The imaging rate is 3 seconds per frame and the play rate is 15 frames per second. The individual locus movements were corrected for the movements of the nuclear centroid for the possible movements of the cell or microscope stage.

**Movie S2. Tracking of IDR2/IDR3 Loci Pair in Cell 2, Related to Figure 4**

The image processing details are the same as described in the movie S1.

## METHODS

### Design RNA Scaffolds for CRISPR-Sirius

To design a stable RNA scaffold accommodating multiple RNA aptamers and compatible for insertion into sgRNA, the aptamers were linked by tandem three way junctions (Filonov et al., 2015). For CRISPR Sirius-8XMS2, we randomized the linker of three way junctions between each MS2 stem loop and made the synonymous mutations of 8XMS2 (Schneider et al., 1992) in the scaffold. To design the variants, we used the consensus sequences as shown below:

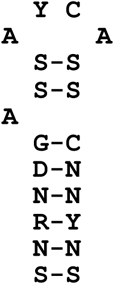

where Y was replaced with C or U, the D was replaced with A, G or U, the S with C or G, R with G or A, and N with any nucleotide. The Sirius-8XMS2 scaffold was designed to avoid a repeating 8-mer in the sequences and to optimize RNA secondary structures (Zuker et al., 2003). The detailed design of Sirius-8XMS2 is shown in Figure S1. The RNA sequence was iteratively evolved by increasing the thresholds (X) for candidate sub-optimal structures. mFold (Zuker et al., 2003) was used to fold the RNA sequence and compute minimum free energy (MFE) and suboptimal free energy (SFE). Initially, all mutable residues were replaced in the base-paring manner shown in the Figure S2 while preserving the A-U or C-G pairs. If the generated sequence contained any repetitive 8-mer, all the mutable residues were mutated again. The sequences were then folded with the initial sub-optimally percentage X=5%. If there was unique structure within the SFE structure, the sequence was fixed and stored. The process was then continued to increase the sub-optimally threshold (X) by 1.0 and the sequence was folded again. Unstable regions were identified if any other structure was predicted within the SFE. Those regions were then marked for further mutation and the process continued until the sub-optimally percentage exceeds 10% or the number of iterations exceeded a given threshold (1000). For CRISPR Sirius-8XPP7, we adapted the three-way junction linkers from CRISPR-Sirius-8XMS2 and used the PP7 aptamer mutants that had the least reduction on the PCP binding (Lim et al., 2002), resulting in the CRISPR Sirius-8XPP7.

### Plasmid Construction

The expression vector for dCas9 (nuclease-dead) from *S. pyogenes* was that originally constructed from pHAGE-TO-DEST (Ma et al., 2015) into which the Heat Stable Antigen (HSA) was inserted at the C-terminus separated by P2A, resulting in pHAGE-TO-dCas9-P2A-HSA. The expression vector for guide RNAs was based on the pLKO.1 lentiviral expression system, in which PUR-P2A-BFP was inserted right after the PGK promoter, with either sgRNA-8XMS2 or sgRNA-8XPP7 inserted immediately after the mouse U6 (mU6) or human U6 (hU6) promoters, resulting pPUR-P2A-BFP-mU6-sgRNA-8XMS2 or pPUR-P2A-BFP-hU6-sgRNA-8XPP7 respectively. The one-step generation of paired guide RNAs was performed by simultaneously subcloning into the cassettes hU6-sgRNA-8XPP7 and mU6-sgRNA-8XMS2 into pPUR-P2A-BFP vector, resulting the dual-guide RNA expression vector pPUR-P2A-BFP-hU6-sgRNA-In-8XPP7-mU6-sgRNA-In-8XMS2, containing the CcdB gene between two BbsI sites in each cassette with different cohesive sites. The details of the cloning strategy were shown in Figure S4. All the dCas9 and guide RNA expression vector reported here will be deposited at Addgene.

### Cell culture, Transfection and Cell Cycle Synchronization

Human osteosarcoma U2OS cells were cultured on 35 mm glass bottom dishes (MatTek) at 37°C in Dulbecco-modified Eagle’s Minimum Essential Medium (DMEM; Life Technologies) containing high glucose and supplemented with 10% (vol/vol) fetal bovine serum. For transfection, typically 200 ng of dCas9 plasmid DNA and 1 µg of plasmid DNA for desired guide RNAs were co-transfected using Lipofectamine 2000 (Life Technologies) and the cells were incubated for another 24-48 hours before imaging. The stable cell line U2OS^IDR2/IDR3^ (see below: “Flow Cytometry and Stable Cell Line Selection”) was synchronized by double Thymidine block based on cell cycle distribution of U2OS (Karanam et al., 2012). Cells were blocked by 2 mM double thymidine for 18 hours, released by rinsing in PBS and then cultured in fresh medium for 9 hours, followed by a second exposure to thymidine for 15 hours (Figure 5A). The cells were then released again and samples taken at 0 hour (early S), 4 hours (mid-late S), 12 hours (early G1) and 17 hours (late G1) respectively.

### Lentivirus Production and Transduction

HEK293T cells were maintained in Iscove’s Modified Dulbecco’s Medium (IMDM; Fisher Scientific) containing high glucose and supplemented with 1% GlutaMAX (Life Technologies), 10% fetal bovine serum (Hycolne FBS, Thermo Scientific) and 1% each penicillin and streptomycin (Life Technologies). 24 hours before transfection, approximately 5x10^5^ cells were seeded in 6-well plates. For each well, 0.5 µg of pCMV-dR8.2 dvpr (Addgene), 0.3 µg of pCMV-VSV-G (Addgene), each constructed to carry HIV LTRs, and 1.5 µg of plasmid containing the gene of interest were co-transfected by using TransIT transfection reagent (Mirus) according to manufacturer’s instructions. After 48 hours, the virus was collected by filtration through a 0.45 µm polyvinylidene fluoride filter (Pall Laboratory). The virus was immediately used or stored at -80 °C. For lentiviral transduction, U2OS cells maintained as described above were transduced by Spinfection in 6-well plates with lentiviral supernatant for 2 days and ∼2x10^5^ cells were combined with 1 ml lentiviral supernatant and centrifuged for 30 minutes at 1200 x *g*.

### Flow Cytometry and Stable Cell Selection

Cells expressing the desired fluorescent Cas9 and/or guide RNA were selected using a FACSAria cell sorter (BD Bioscience) equipped with 405, 488, 561 and 640 nm excitation lasers, and the emission signals were detected by using filters at 450/50 nm (wavelength/bandwidth) for the Brilliant Violet 421-conjugated anti-mouse CD24 antibody (BioLegend) staining of the HSA, 530/30 nm for PCP-GFP and 582/15 nm for MCP-HaloTag stained with HaloTag-JF549. For the sorting of dCas9 signals, 1 µl of the Brilliant Violet 421-conjugated anti-mouse CD24 antibody was added in a 100 µl cell solution for 30 minutes before FACS. For sorting of MCP-HaloTag, HaloTag-JF549 was added to the cells at 2 nM 18-24 hours before sorting. Single cells were sorted into single wells of 96-well plates containing 1% GlutaMAX, 20 % fetal bovine serum and 1% penicillin and streptomycin in chilled DMEM medium. Positive clones of U2OS^dCas9-HSA/PCP-GFP/MCP-HaloTag^ were selected from 96-well plates 10 days later. To generate stable cell lines in which the IDR2/IDR3 locus pair was durably labeled, the U2OS^dCas9-HSA/PCP-GFP/MCP-HaloTag^ cell line was transduced for 48 hours by lenti-virius for PUR-P2A-BFP-mU6-IDR2-sgRNA-8XMS2-hU6-IDR3-sgRNA-8XPP7-IDR3 for 48 hours. Cells were then selected on puromycin for 3-5 days before sorting for BFP, using filters at 405 nm excitation and 450/50 nm emission. The resulting cell lines was simply named U2OS^IDR2/IDR3^.

### Fluorescence Microscopy

A Leica DMIRB microscope was equipped with an EMCCD camera (Andor iXon-897), mounted with a 2x magnification adapter and 100x oil objective lens (NA 1.4), and resulting in a total 200x magnification equal to a pixel size of 80 nm in the images was used. The microscope stage incubation chamber was maintained at 37 °C in HEPES-buffered DMEM with 10% FBS. GFP was excited with an excitation filter at 470/28 nm (Semrock) and its emission was collected using an emission filter at 512/23 nm (Semrock). HaloTag-JF549 was excited at 556/20 nm (Semrock) and its emission was collected in a 630/91 nm channel. Imaging data were acquired by MetaMorph acquisition software (Molecular Devices). Image size was adjusted to show individual nuclei and intensity thresholds were set on the basis of the ratios between nuclear focal signals to background nucleoplasmic fluorescence. To detect loci numbers in Figure 3B, maximum intensity projection of Z-series images was performed. To quantify the spatial distance or track the dynamics, only pairs of loci lying in the same foci plane were analyzed.

## QUANTIFICATION AND STATISTICAL ANALYSIS

### Mining Chromosome-Specific Repeats for the Human Genome

Human reference genome (assembly GRC h37/hg19) (genome.ucsc.edu) was analyzed to find target regions and design gRNAs. Bioinformatics tool Tandem Repeat Finder (Benson et al., 1999) was used to identify tandem repeats with repeats period length smaller or equal to 2000 bp in the human genome. Bioinformatics tool Jellyfish (Marçais et al., 2011) was used to identify tandem repeats with repeat length longer than 2000 bp in human genome. Jellyfish was used to search for the 15-mers in the identified repeat regions. All the tandem repeat regions with more than 5 non-overlapping copies of one 15-mer were selected. The non-overlapping repetitive 15-mers with CRISPR PAM sequences ending with NGG or starting with CCN were examined for their specificity. The 15-mers that had more than 20% of copies within in other 50 kb regions were discarded. The 15 mers containing “TTTT” or ending with “TNGG” were filtered out due to the potential pre-termination on sgRNA expression under the U6 promoter (Chen et al., 2013). The distribution of unique repeats in human genome was shown in Figure 1 and chromosome 19-specific unique repeats identified by the above-mentioned bioinformatics pipeline was shown in Figure S2. The copy numbers shown in the figures were defined as the maximal non-overlapping target sites from a single sgRNA in the region.

### Imaging Processing of Pairs of Loci Tracking

The images were analyzed by the *Fiji* (http://fiji.dc/Fiji) and *Mathematica* (Wolfram) software. Images from the green and red channels were registered by using 0.1 µm coverglass-absorbed TetraSpeck fluorescent microsphere (Invitrogen) as a standard sample. Intensity quantification in Figure 2F was performed as following

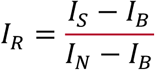

where I_R_ is the intensity ratio between the labeled FBN3 loci (I_S_) and nucleoplasm (I_N_). each with is the background fluorescence intensity (I_B_) from a dark region in the same image having been subtracted. In live cell tracking, the specific genomic loci signals were identified and tracked by using the TrackMate plugin (Tinevez et al., 2016). The cell movement was corrected by tracking the centroid of the nucleus over time. Detailed calculation of the mean-square displacement (MSD), relative spatial distance (D), gyration radius of trajectory radius (R) and relative angle (θ_***R***_) are described in following section.

### Quantification of Distance and Movement from Locus Pairs

Let ***p***(*t*) be the position vector of a locus at time t. The trajectory of the locus is defined by a time series of positions {***p***_0_, ***p***_***1***_,…,***p***_***n***_} where ***p***_***k***_:=***p***(*t*_*k*_*), t*_*k+1*_=*t*_*k*_*+Δt,* 0=*t*_*0*_*<t*_*1*_*,…,<t*_*n*_ and *Δt* is a fixed time interval between two successive frames. Given a trajectory {***p***_0_, ***p***_***1***_,…,***p***_***n***_}, the associated mean-squared displacement (MSD) of lag time *kΔt* is given by

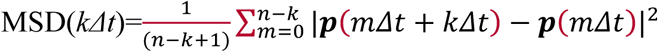

Given trajectories {***p***_***A***_(*t*_*0*_), ***p***_***A***_(*t*_*1*_),…,***p***_***A***_(*t*_*n*_)} and {***p***_***B***_(*t*_*0*_), ***p***_***B***_(*t*_*1*_),…,***p***_***B***_(*t*_*n*_)} of two loci A and B, the relative and centroid trajectories are defined by {***p***_***AB***_(*t*_*0*_), ***p***_***AB***_(*t*_*1*_),…,***p***_***AB***_(*t*_*n*_)} and {***p***_***C***_(*t*_*0*_), ***p***_***C***_(*t*_*1*_),…,***p***_***C***_(*t*_*n*_)}, respectively where ***p***_***A/B***_(*t*_*k*_):= ***p***_***B***_(*t*_*k*_)-***p***_***A***_(*t*_*k*_) with the length D_***AB***_(*t*_*k*_):=|***p***_***AB***_(*t*_*k*_)| and 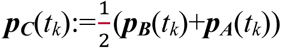 for *k=0, 1,…,n*. Their associated MSDs are denoted by MSD_***A/B***_ and MSD_***C***_ respectively. All MSD curves were fitted by the power law equation, 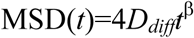 where *D*_*diff*_ is the diffusion constant. All MSDs possessed exponents 0< *β*<1. In addition to the plots in the figures, all the values are listed in Table S1.

We also defined a time series of the relative angle {θ_***R***_(*t*_*1*_), θ_***R***_(*t*_*2*_),…, θ_***R***_(*t*_*n*_)} for the relative trajectory where θ_***R***_(*t*_*m*_) is defined by the angle swept from the vector ***p***_***AB***_(*t*_*m-1*_) to the vector ***p***_***AB***_(*t*_*m*_) for *m=1, 2,…,n*. The angle was calculated by

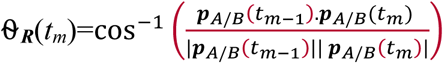

where −180≤ θ_*R*_ (*t*_*m*_) ≤ 180° with the sign reflecting the orientation of the angle, positive for counterclockwise and negative for clockwise.

In addition to MSD, we also introduce a quantity, named the trajectory radius, to measure the mobility of a locus or locus pair. This trajectory radius is defined as the gyration radius of the points collected from all steps on a trajectory and used to measure the range of the area covered by the trajectory. More precisely, given trajectory {***p***_***S***_(*t*_*0*_), ***p***_***S***_(*t*_*1*_),…,***p***_***S***_(*t*_*n*_)}, ***S***=***A, B, A/B, C***, the trajectory radius Rsof trajectory ***S*** is defined by

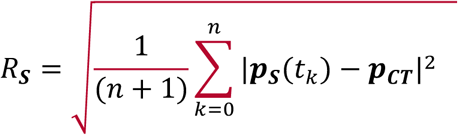

Where 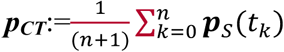 is the geometric center of the positions {***p***_***S***_(*t*_*0*_), ***p***_***S***_(*t*_*1*_),…,***p***_***S***_(*t*_*n*_)}.*α* denotes the scaling exponent of the power law relationship, mean spatial distance *∝* (genomic distance) *α* and is determined by fitting the power law relationship with experimental data.

All these analyses including the cumulative probability of loci pairs’ spatial distance were performed by *Mathematica* and graphs were generated by *OriginPro* (OriginLab). All box plots were generated using the default setting of the *OriginPro*, box spans from first to last quartiles and whisker length is determined by the outermost data point that falls within upper inner and lower inner fence (a coefficient =1.5), except the middle line represents the mean value.

